# An aorta-to-insulin producing cell sensor for internal state in *Drosophila*

**DOI:** 10.64898/2026.06.17.732871

**Authors:** Anton Miroschnikow, Andreas Schoofs, Philipp Schlegel, Damian Demarest, Marei Freitag, Albert Cardona, Constantin Pape, Michael J Pankratz

## Abstract

The cellular circuits through which the internal state of an organism is sensed and relayed to the brain, and through which they bring about appropriate physiological and behavioral responses, remain largely unelucidated. Using whole-animal electron microscopy reconstruction of a Drosophila larva, we previously identified an Aorta sensory organ whose somata lie on the periphery of the brain, with dendritic arborizations in the wall of the anterior aorta, and whose axons project to the protocerebrum and synapse onto the insulin-producing cells (IPCs). Here we characterize the full anatomy and connectivity of this sensory organ. We show that the aorta sensory neurons make their strongest direct synaptic contacts onto IPCs and the neurosecretory cells producing Dromyosuppressin (DMS), with only weaker direct contacts to diuretic hormone 44 (DH44). They also make direct contacts to the enteric serotonergic neurons modulating swallowing (Se0ens), and to two single mushroom body neurons, DAN-j1 and MBON-d3, of the learning and memory center of Drosophila. Calcium imaging shows that the neurons are activated by fructose, and optogenetic manipulation reveals that their activity influences peptide content in IPCs, DH44 and DMS cells. Polysynaptic paths to the brain run through two morphologically distinct interneuron families. One, formed by the Hugin protocerebral neurons, targets the neuroendocrine system and is associated with bitter taste and pathogen induced feeding suppression. The other, formed by the BAmas projection neurons, targets dopaminergic input neurons and output neurons of the mushroom body. The Aorta sensory pathway therefore couples internal hemolymph state to two parallel central systems — endocrine state control and mushroom body-associated value updating — and provides a candidate substrate for a metabolic memory through which post-ingestive state is fed back into future feeding choice.

## INTRODUCTION

Animals must keep track of their own internal nutritional state and turn that information into appropriate metabolic, endocrine, and behavioral response. Insulin sits at the centre of this regulation, and its role is conserved from worms to mammals. In vertebrates it is made by the pancreatic β-cells and released into the blood, where it acts on target tissues through the insulin receptor (Steiner 1969; Saltiel and Kahn 2001). Across phyla, insulin and insulin-like peptide signaling systems regulate metabolism, growth, and lifespan, with extensive work identifying the nutrient cues that control their activity and the downstream pathways through which they act (Brogiolo et al. 2001; Garofalo 2002; Kenyon 2010). In Drosophila, systemic insulin signal originates from a small group of insulin-producing cells (IPCs) in the pars intercerebralis (PI), alongside neurosecretory cells producing diuretic hormone 44 (DH44) and Dromyosuppressin (DMS). (Rulifson et al. 2002; Cao and Brown 2001; Nässel and Vanden Broeck 2016). How these cells are regulated, and what happens when their activity is compromised, has been studied in considerable detail (Suzawa and Bland 2023). Some of the routes that link nutritional state to this central endocrine output are now known. DH44 neurons, for instance, are activated by nutritive sugars and dietary amino acids and are needed for post-ingestive food choice (Yang et al. 2018; Dus et al. 2011; Oh et al. 2019), and the fat-body- and gut-derived peptide CCHamide-2 acts on its receptor on the IPCs to couple peripheral nutrient state to insulin release and memory consolidation (Sano et al. 2015; Ren et al. 2015; Yamagata et al. 2022).

The complementary question is how the central nervous system itself learns about the body’s metabolic state. In vertebrates, circumventricular organs and vagal afferents function in this capacity. Circumventricular organs are small brain regions in which the blood-brain barrier is relaxed, allowing neurons to sample circulating peptides, nutrients, and hormones directly from the bloodstream (Ganong 2000; Fry et al. 2007). Through them, insulin, leptin, ghrelin, glucose, and gut-derived peptides gain access to hypothalamic and brainstem circuits that govern feeding and energy balance (Johnson and Gross 1993; McKinley et al. 2003; Miyata 2022; Schwartz et al. 2000; Brüning and Fenselau 2023). In parallel, the brain receives internal-state information through visceral afferents of the vagus nerve, whose endings monitor gastric distension, nutrient absorption, and circulating signals (Berthoud and Neuhuber 2019; Williams et al. 2025; Han et al. 2018). Insects face the same problem: how does the CNS gain timely access to circulating internal-state information? One possibility is that information from the hemolymph reaches the brain through the blood-brain barrier and associated glial cells (Hertenstein et al. 2021). Such routes may support broad metabolic monitoring, but they are unlikely to provide the fast and anatomically specific access needed to couple circulating state information directly to defined neuroendocrine circuits. Yet whether insects possess dedicated hemolymph-facing sensory cells that perform this function has remained unclear. Drosophila larvae provide a particularly useful setting to address this question, because feeding, growth, and systemic endocrine state are tightly coupled and the full circuit can be reconstructed at synaptic resolution. The missing element is anatomical: a sensory interface with access to the hemolymph on one side and direct projections to endocrine circuits in the brain on the other.

In this study, we provide a complete morphological and connectivity analysis of the aorta-sensory neurons in the larva. These neurons were encountered several times before they were recognised. They first appeared in the hugin neurosecretory cell connectome as unidentified sensory afferents contacting huginPC neurons and median neurosecretory cells, including the IPCs (Schlegel et al. 2016). The larval feeding connectome then placed this projection within a broader SEZ architecture linking sensory inputs to feeding motor neurons, serotonergic output neurons, and PI neuroendocrine cells (Miroschnikow et al. 2018), while the neuroendocrine connectome further resolved how sensory and interneuronal pathways converge onto distinct endocrine outputs (Hückesfeld et al. 2021). The peripheral origin of the projection, however, remained unresolved until Schoofs et al. annotated the corresponding cells as aorta-sensory neurons and localized their somata, linking them to the aortic region (Hückesfeld et al. 2021; Schoofs et al. 2024). Their peripheral morphology, dendritic target site, and full downstream connectivity remained unknown.

Here we resolve that anatomy and reconstruct the full synaptic input and output of the larval aorta-sensory neurons. We show how this sensory system links the hemolymph compartment to central endocrine, feeding, and memory-related circuits, including insulin-producing cells and mushroom body dopaminergic neurons. On this basis, we propose that the aorta-sensory organ provides a route through which internal metabolic state can be read out together with memory systems to coordinate motor and endocrine output during feeding. The corresponding cells are also present in the adult brain, and we concisely relate the larval and adult anatomy, but our focus is the larva, where the entire circuit can be reconstructed at synaptic resolution.

## RESULTS

### Anatomy of the aorta sensory organ: peripheral and central projections

Using a combination of whole-animal EM reconstruction and light-microscopic analysis, we elucidated the morphology of the larval aorta-sensory system in the context of other major organ systems (**Figure 1A**). The aortic funnel forms a widened anterior end of the aorta above the esophageal region. The dendritic field of the aorta-sensory neurons is embedded in the brain-facing funnel wall and exposed to the hemolymph compartment (**Figure 1B**). This region lies close to the esophageal mechanosensory neurons and ring muscle motor innervation described earlier by our lab (Schoofs et al. 2024), but this proximity reflects the geometry of the funnel rather than projections wrapping around the esophageal wall.

**Figure 1.**
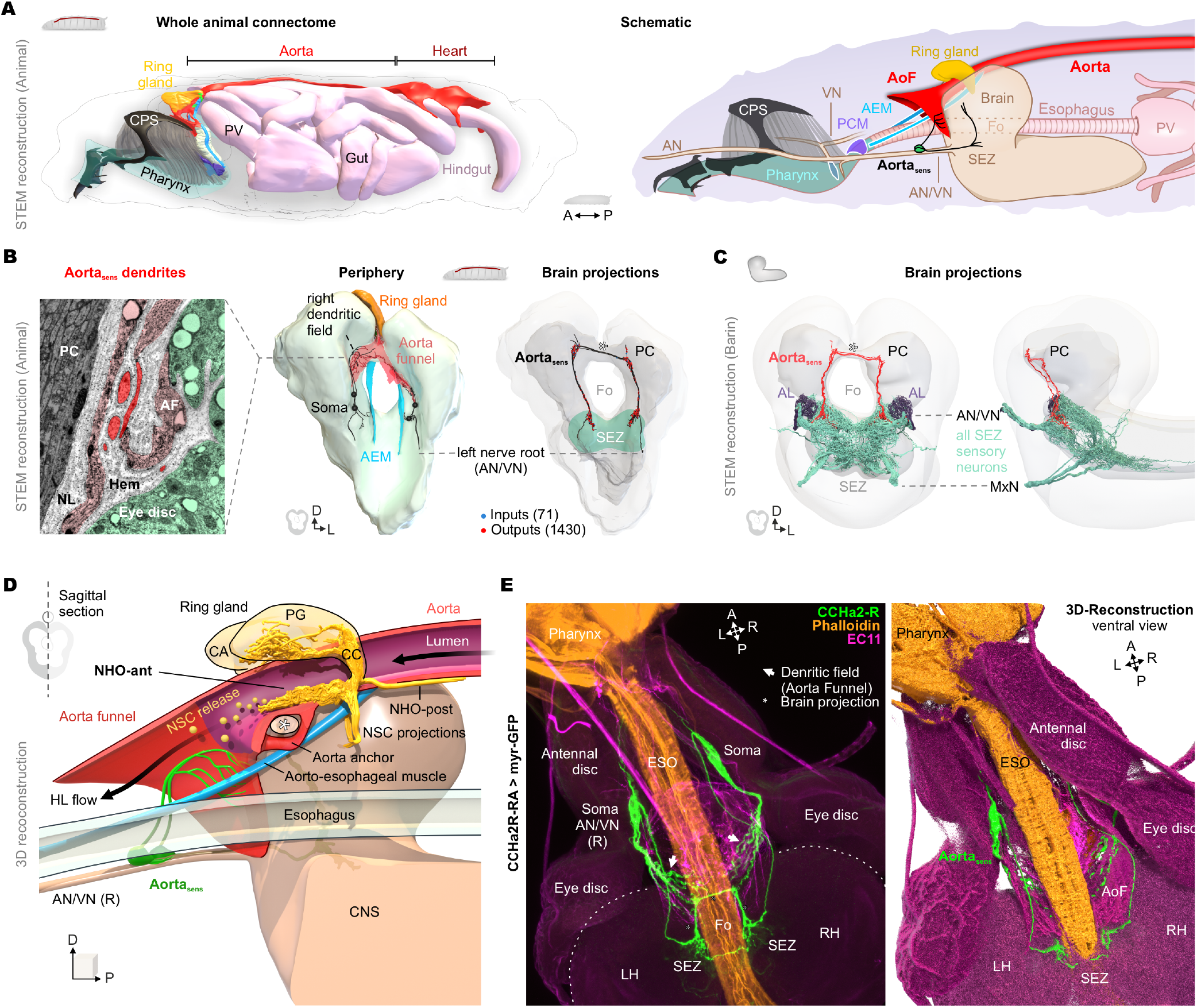
Aorta sensory neurons form a sparse peripheral-to-brain interoceptive pathway. **(A)** 3D reconstruction of the whole-larva EM volume *Igor*, showing the CNS, pharynx, digestive tract, heart/aorta, aortic funnel, and ring gland. The right panel shows a simplified schematic of the same anatomical arrangement. The whole-animal EM volume provides single-cell and synapse-level anatomical ground truth linking central projections to peripheral tissues. **(B)** Reconstruction of aorta sensory neurons in *Igor*. Central projections terminate in the brain, whereas peripheral processes extend into the aortic funnel. Representative EM sections show dendritic profiles embedded in the aortic funnel region and positioned at a hemolymph-facing interface. **(C)** Reconstruction of the same neuron class in the larval brain EM volume. Among sensory neurons projecting into the brain, aorta sensory neurons represent a sparse internal sensory pathway with projections to the protocerebrum. **(D)** 3D model of the brain-associated aorta complex. A sagittal section through the aorta, ring gland, brain, and esophagus reveals the anterior aortic interface. Neuroendocrine projections from the brain enter the aorta, contribute to the aortic wall, and form the neurohemal organ, where neuropeptides are released into the aorta-associated hemolymph space. Aorta sensory dendrites are positioned immediately posterior to this region within the aortic funnel wall. The aorta is anchored at the brain midline by an aortic anchor, and a muscle connects the neurohemal organ to the pharynx–esophagus transition. **(E)** CCHa2R-RA-Gal4 crossed to UAS-myr::GFP labels a sparse neuronal population matching the EM-reconstructed aorta sensory neurons. EC11 (anti-Pericardin) staining marks the cardiac/aortic extracellular matrix. The right panel shows a 3D reconstruction of the confocal data generated in Fiji and rendered in Blender.

The somata of the aorta-sensory neurons reside on the compound antennal/pharyngeal/vagal nerve shortly before its entry into the brain. From there, their axons follow this nerve into the SEZ, ascend into the protocerebrum, cross the midline through the brain commissures, and descend contralaterally back to the SEZ (**Figure 1B,C**). Comparison with the complete SEZ sensory neuron population shows that the aorta-sensory neurons are the only identified sensory neurons in this set that extend beyond the SEZ into the protocerebrum (**Figure 1C**).

A detailed reconstruction of the aortic region, with the aorta virtually opened by a sagittal cut, revealed the relative positions of the neurohemal organ and the aorta-sensory dendrites (**Figure 1D**). The same reconstruction also revealed an aorto-esophageal muscle (AEM), described here for the first time (**Figure 1A, D**). We did not detect motor neuron innervation of this muscle. The AEM attaches anteriorly to a small muscle-free zone of the esophagus immediately posterior to the pharyngeal constrictor muscle (PCM), and posteriorly to the aortic wall just behind the corpora cardiaca (CC). In addition to the anterior neurohemal organ (NHO), which contains release sites inside the aorta, we identified a previously undescribed posterior extension, NHO-post. This posterior component is formed by IPC, DMS, and DH44 projections that run along the surface of the aorta, using the vessel as an anatomical scaffold. The aorta itself is held in place by an aortic anchor surrounding the commissural region between the two brain hemispheres.

A sparse CCHa2R-RA-Gal4 line that labels the aorta-sensory neurons allowed us to validate this organization at the light-microscopic level (**Figure 1E**). Together with the nearby PI neurosecretory projections into the aorta and the monosynaptic input of aorta-sensory neurons onto IPCs, DMS, and DH44 neurons, this arrangement defines a direct anatomical loop between the hemolymph-facing aortic interface and central neurosecretory circuits

### Insulin-producing cells are the major monosynaptic targets of the Aorta sensory neurons

We next identified the direct postsynaptic partners of the aorta-sensory neurons in the CNS-only (“Seymour”) EM volume (**Figure 2A**). Fourteen percent of their total synaptic output targeted functionally annotated neurons, including neuroendocrine cells, serotonergic modulatory output neurons, and mushroom body–related interneurons (**Figure 2B,C**). Within this set, IPCs and DMS neurons were the strongest neuroendocrine targets, whereas DH44 and eclosion hormone (EH) neurons received weaker input. The sparse mushroom body–related contacts are notable because they place hemolymph-associated sensory input in direct reach of dopaminergic and memory-related circuitry.

**Figure 2.**
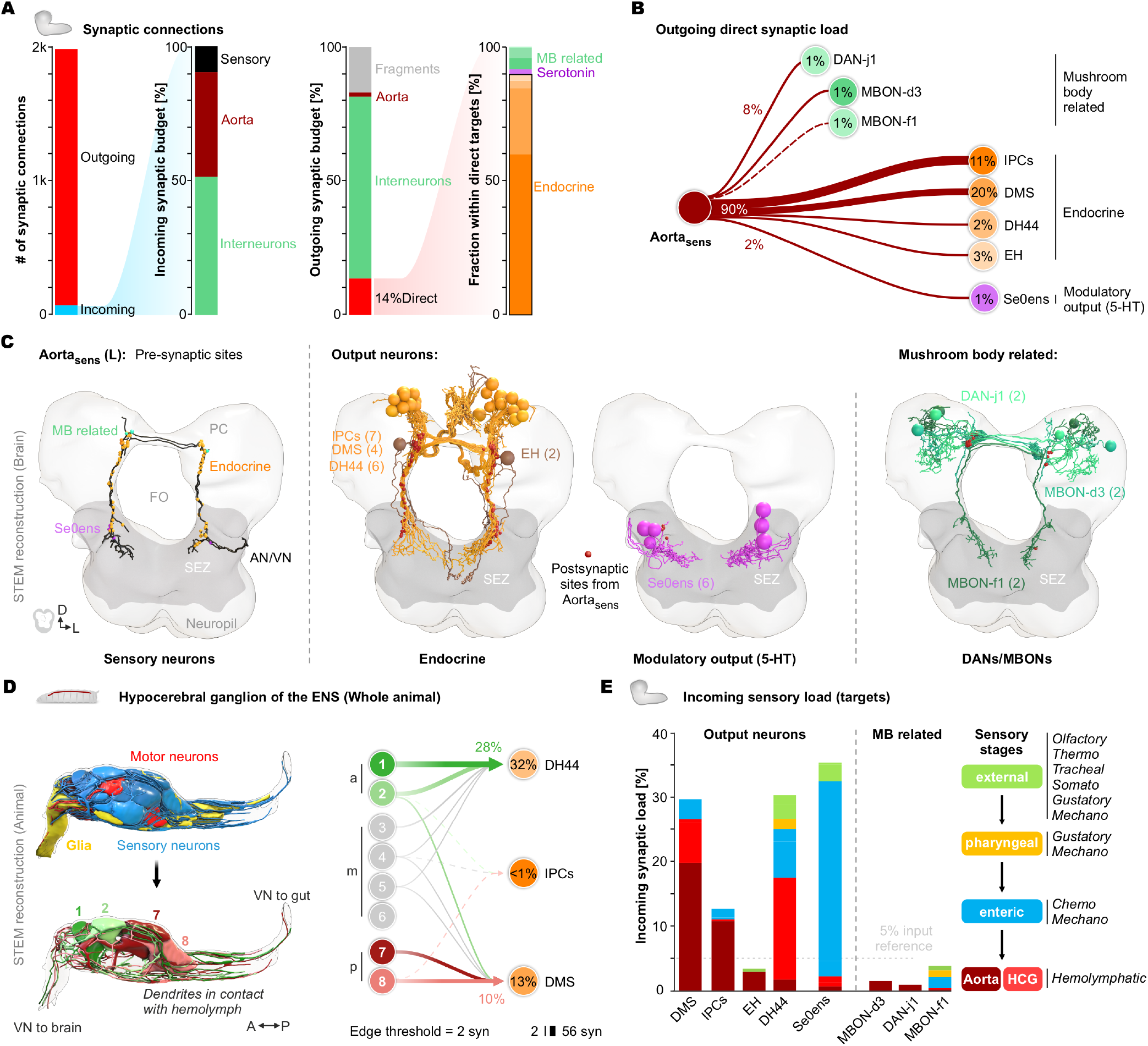
Aorta sensory neurons directly connect internal hemolymph-associated sensory input to neuroendocrine, modulatory, and mushroom body–related circuits. **(A)** Synaptic input and output architecture of aorta sensory neurons. Left half, total incoming and outgoing synapses, and normalized composition of presynaptic inputs, highlighting feedback-like inputs onto aorta sensory neurons. Right half, normalized composition of direct postsynaptic targets. Aorta sensory neurons allocate 14% of their direct output to neuroendocrine neurons, with additional connections to serotonergic modulatory output neurons and mushroom body–related neurons. **(B)** Node graph of direct postsynaptic targets. Edge weights indicate the normalized fraction of total outgoing aorta sensory synapses. Percentages within target nodes indicate the fraction of each target’s total input contributed by aorta sensory neurons. **(C)** Left most, unilateral EM reconstruction of left-sided aorta sensory neurons, showing morphology and synaptic sites onto the targets. Synapses are marked as spheres. Right, reconstructions of major target classes, including neuroendocrine neurons, Se0ens serotonergic output neurons, the PAM-cluster dopaminergic neuron DAN-j1, and two mushroom body output neurons. **(D)** Volumetric ultrastructural reconstruction of the hypocerebral ganglion (HCG) from the whole-animal connectome. The HCG contains five esophageal motor neurons and eight sensory neurons. Glia ensheath neuronal processes up to the soma region, whereas somata and dendrites remain exposed to the hemolymph. Anterior HCG sensory neurons 1 and 2 (HCGa) provide major monosynaptic input to DH44 neurons, whereas posterior sensory neurons 7 and 8 (HCGp) provide major monosynaptic input to DMS neurons, as summarized in the node graph on the right. **(E)** Sensory input load of direct aorta sensory neuron targets compared with other sensory classes. Direct targets integrate external, pharyngeal, enteric, and internal sensory inputs. DH44 neurons are dominated by HCG input but also receive broad multisensory input. Right, sensory-stage scheme classifying inputs from external to pharyngeal, enteric, and internal hemolymph-associated pathways, including aorta sensory and HCG neurons.

We then compared this output with other internal sensory pathways. The hypocerebral ganglion (HCG) neurons had previously been reconstructed (Schoofs et al. 2024); here, ultrastructural analysis showed that their somata and dendrites are exposed to the hemolymph, whereas ENS axons are ensheathed by glia from the brain up to the HCG soma region (**Figure 2D**). This identifies the HCG as a second hemolymph-associated sensory site. Combining their anatomy with their characteristic connectivity in the whole-animal volume—particularly their dominant sensory input to specific neuroendocrine targets— allowed us to assign the corresponding HCG neurons in the brain volume and place them within the broader sensory input architecture. Anterior HCG sensory neurons provided strong monosynaptic input to DH44 neurons, whereas posterior HCG sensory neurons provided prominent input to DMS neurons. Thus, aorta-sensory neurons provide strong direct access to selected neuroendocrine cells, while each target integrates a distinct sensory profile across aorta-sensory and HCG sensory neurons **(Figure 2E**).

### Aorta-sensory neurons respond to fructose and regulate peptide content in median neurosecretory cells

This anatomical organization raised the question of whether aorta-sensory neurons are functionally linked to nutrient state and whether their activity can influence the neuroendocrine cells they contact. Aorta sensory neurons express several genes associated with nutrient and metabolic function, including the fructose receptor Gr43a (Miroschnikow et al. 2018; Schoofs et al. 2024; Miyamoto et al. 2012). We therefore tested whether these neurons respond to circulating sugars. Live calcium imaging of GCaMP6f-expressing Aorta sensory neurons showed a selective response to 50 mM fructose, whereas glucose did not evoke significant activation and trehalose reduced activity (**Figure 3A,B**).

**Figure 3.**
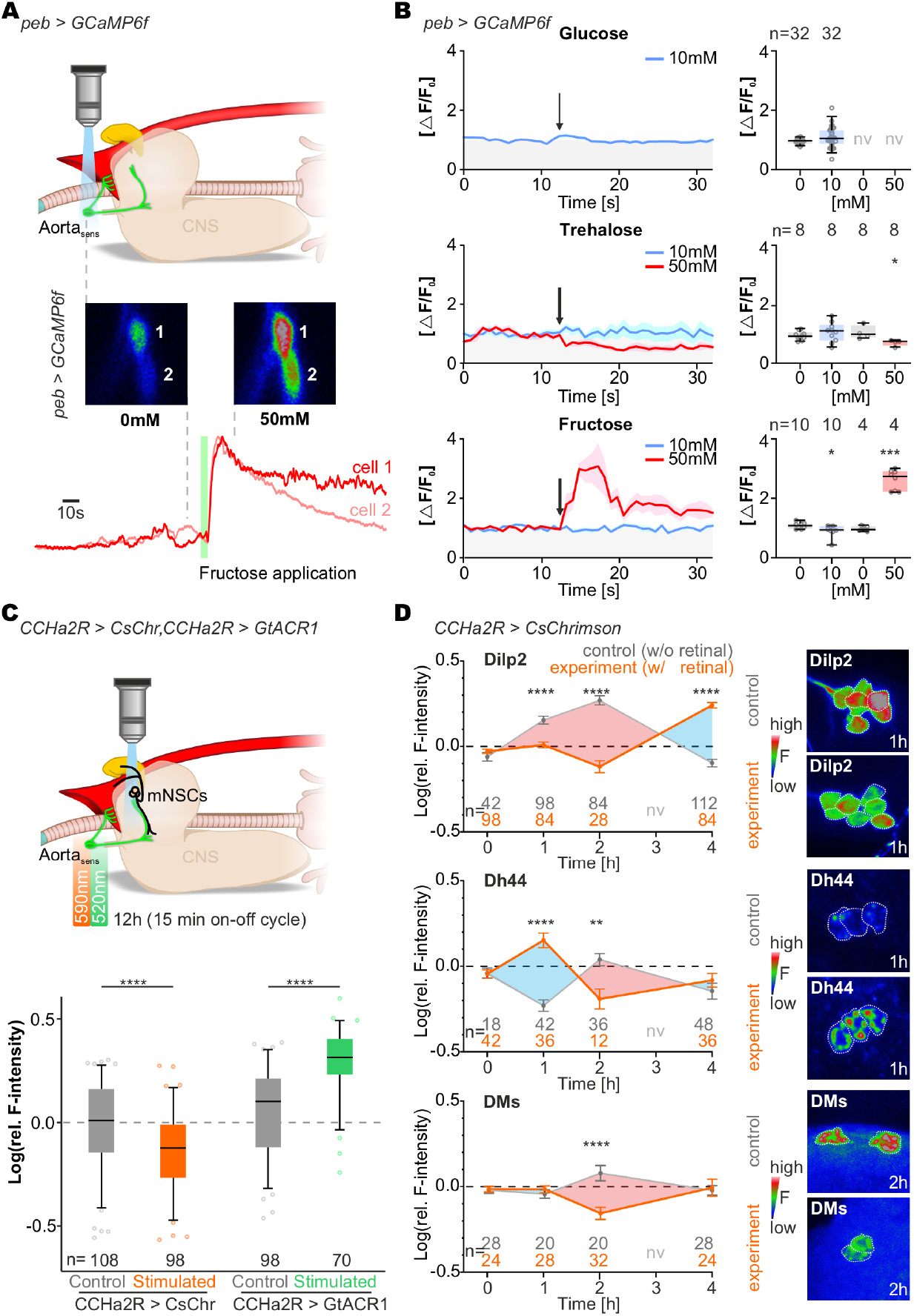
Aorta sensory neurons respond to fructose and regulate peptide content in median neurosecretory cells. (A) Live calcium imaging of aorta sensory neurons expressing GCaMP6f under control of *pebbled*-Gal4. Example images show two left-sided aorta sensory neurons before and after application of control solution or 50 mM fructose; traces show corresponding calcium responses. **(B)** Quantification of responses to glucose, trehalose, and fructose. Aorta sensory neurons showed sugar-evoked activity selectively to 50 mM fructose, whereas 50 mM trehalose reduced activity. Statistics: Mann–Whitney U test. **(C)** Bidirectional optogenetic manipulation of aorta sensory neurons using CCHa2R-RA-Gal4 to drive CsChrimson or GtACR1. Larvae were stimulated for 12 h with 15-min light-on/light-off cycles using 590 nm light for CsChrimson activation or 520 nm light for GtACR1 inhibition. Dilp2 immunofluorescence decreased after CsChrimson-mediated activation, consistent with peptide depletion and putative release, and increased after GtACR1-mediated inhibition, consistent with peptide accumulation. **(D)** Quantitative peptide immunostaining after continuous 590 nm activation of aorta sensory neurons. Larvae were collected after 0, 1, 2, and 4 h and stained for Dilp2, DH44, or DMS. Dilp2 showed progressive depletion up to 2 h, followed by peptide re-accumulation at 4 h. DH44 showed a fluctuating response across time points, with alternating decreases and increases in immunofluorescence. DMS showed a significant increase in inferred peptide depletion only after 2 h. Representative pseudocolored antibody stainings for control and experimental conditions are shown on the right. Statistics: Mann–Whitney U test.

We next asked whether the activity of Aorta sensory neurons is sufficient to alter peptide content in their neuroendocrine targets. Using CCHa2R-RA-Gal4, we bidirectionally manipulated Aorta sensory neurons with CsChrimson or GtACR1 for 12 h and quantified Dilp2 immunofluorescence in IPCs (**Figure 3C**). Activation decreased Dilp2 levels, consistent with peptide depletion and putative release, whereas inhibition increased Dilp2 levels, consistent with peptide accumulation. To resolve the temporal structure of this effect, we then performed an activation time course over 1, 2, and 4 h and extended the analysis to DMS and DH44 neurons (**Figure 3D**). Across all three peptide systems, activation produced temporal profiles that were complementary to the corresponding controls. Dilp2 showed progressive depletion up to 2 h, followed by re-accumulation at 4 h. DH44 displayed alternating decreases and increases in peptide immunofluorescence across the time course, whereas DMS changed more selectively, with significant inferred peptide depletion after 2 h. The physiological meaning of these dynamics remains unresolved, but the target-specific responses are consistent with a circuit architecture in which the IPCs and DH44 neurons are targeted distinctly by aorta-sensory and HCG sensory neurons.

### Aorta sensory neurons in adults

We next asked whether the Aorta sensory pathway is also present in the adult Drosophila brain and identified the corresponding neuron class, CB0991, in the adult FAFB EM volume (Dorkenwald et al. 2024; Schlegel et al. 2024; Zheng et al. 2018). CB0991 had previously been linked to CCHa2R-RA-Gal4 expression and neuroendocrine control of sugar and water ingestion, but was interpreted as an interneuron population rather than as a sensory pathway associated with the aorta (González Segarra et al. 2023). Guided by the larval anatomy and CCHa2R-RA-Gal4 expression, we therefore re-examined CB0991 in FAFB.

Adult Aorta sensory neurons showed direct output to neuroendocrine cells, which accounted for 15% of their total synaptic output (**Figure 4A,B**). As in the larva, IPCs were among the strongest neuroendocrine targets. A notable difference was a weaker connection to DMS neurons in the adult compared with the larval circuit. CCHa2R-RA-Gal4 expression further supported the anatomical correspondence between the larval and adult neuron classes (**Figure 4C,D**). Although the adult anatomy differs substantially from the larva, several core features are retained: an aortic funnel at the anterior end of the aorta, dendritic processes positioned near this funnel, and central projections that provide synaptic output to IPCs. In the adult, the aortic funnel is associated with a peripheral muscle support site related to the antennal-heart system, likely corresponding to M16 described in classical and recent anatomical work (Demerec 1950; Kay et al. 2021). EM reconstruction of CB0991 confirmed this organization and showed dendritic processes extending along the posterior brain surface toward the foramen, near the aortic funnel and antennal-heart-associated structures (**Figure 4E**).

**Figure 4.**
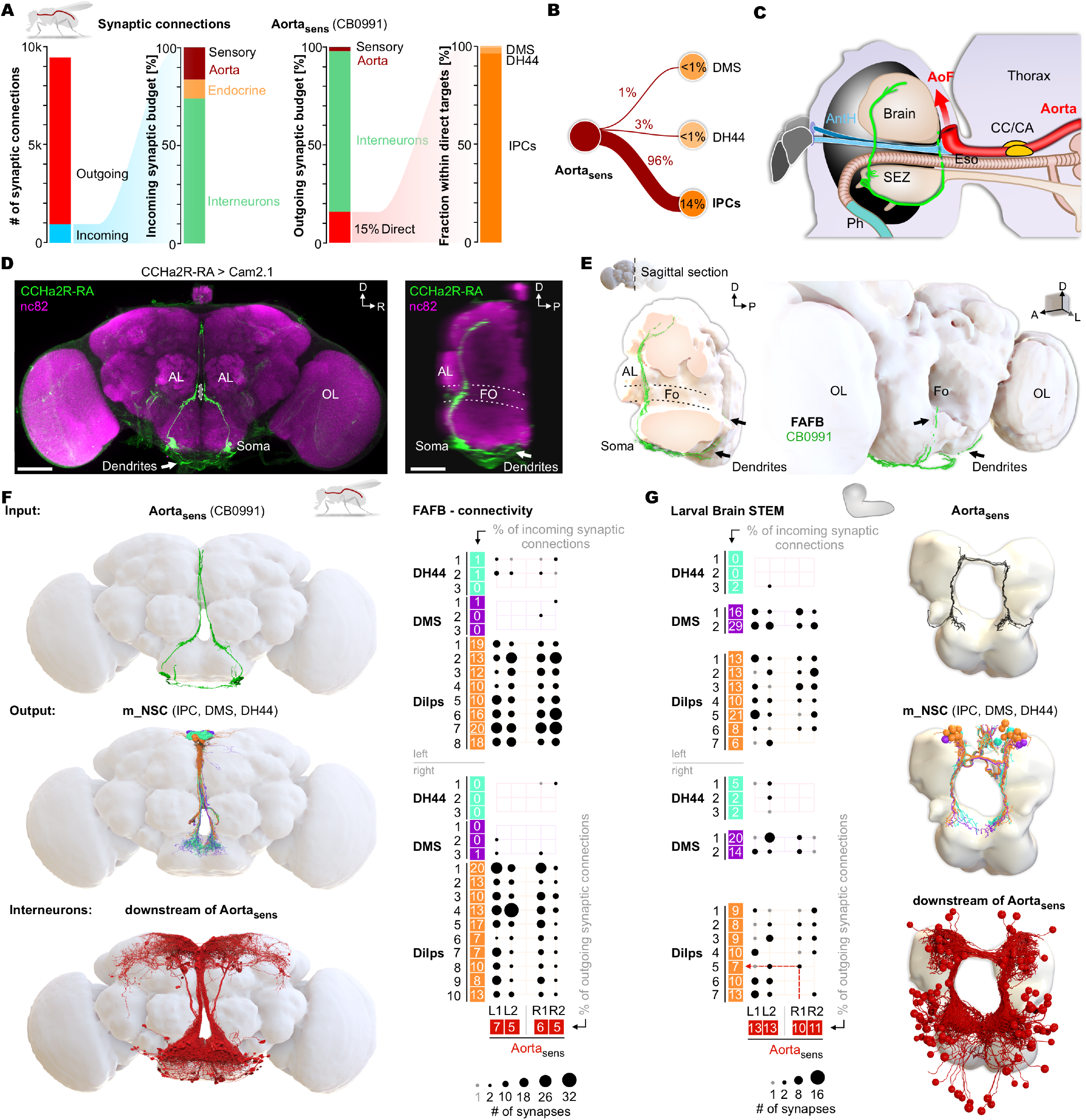
Conserved adult and larval aorta sensory pathways target neuroendocrine circuits. **(A)** Synaptic input and output architecture of adult aorta sensory neurons. Left half, total incoming and outgoing synapses, and normalized composition of presynaptic inputs, including feedback-like inputs onto aorta sensory neurons. Right half, normalized composition of direct postsynaptic targets, showing prominent output to neuroendocrine neurons that include 15% of direct synapses. **(B)** Node graph of direct postsynaptic targets of adult aorta sensory neurons. Edge weights indicate the normalized fraction of total outgoing aorta sensory synapses. Percentages within nodes indicate the fraction of each target’s total synaptic input contributed by aorta sensory neurons. (**C**) Scheme of adult aorta sensory neuron projections and aorta/aorta funnel architecture. **(D)** CCHa2R-RA-Gal4 expression in the adult brain, shown in frontal and lateral views. In the lateral view, aorta sensory neuron processes extend along the ventral brain surface beneath the SEZ toward the neck connective. **(E)** EM reconstruction of the same neuron class in the adult FAFB volume. A sagittal cut through the brain reveals the position and trajectory of the aorta sensory neurons, supporting the lateral organization observed in C. Posterior view shows that the dendritic processes of the aorta sensory neurons, corresponding to CB0991 in FAFB, extend along the posterior brain surface toward the foramen, near the aortic funnel and antennal-heart-associated structures. **(F)** Adult FAFB reconstructions of aorta sensory neurons/CB0991, their direct monosynaptic neurosecretory targets, and downstream interneurons. The connectivity diagram on the right shows single-cell monosynaptic connections from aorta sensory neurons to DH44-, DMS-, and Dilp-positive median neurosecretory cells. **(G)** Corresponding larval connectivity and reconstructions from the larval brain EM volume. As in F, aorta sensory neurons, their direct neuroendocrine targets, and downstream neurons are shown, highlighting conserved connectivity to DH44, DMS, and Dilp neurosecretory populations across developmental stages.

Extending the connectivity analysis to additional downstream targets further highlighted similarities between larval and adult Aorta sensory pathways (**Figure 4F,G**). In both stages, Aorta sensory neurons contact Dilp-, DMS-, and DH44-positive median neurosecretory populations, although the relative strength of these connections differs between larva and adult. Thus, despite major developmental changes in anatomy, the Aorta sensory pathway retains a conserved association with neuroendocrine circuits across life stages. Recently, additional pathways involving the aorta-sensory neurons have been identified in the adult whole CNS connectome, and it would be interesting to see how these circuits are organized in the larva (Bates et al. 2026).

### Aorta sensory neurons distribute hemolymph-associated input through distinct interneuron relays

Previous connectomic analyses showed that sensory information reaches feeding motor, serotonergic, and neuroendocrine outputs through multiple parallel interneuron paths (Miroschnikow et al. 2018; Hückesfeld et al. 2021). We therefore asked whether Aorta sensory neurons use a similar relay architecture to distribute hemolymph-associated input beyond their direct monosynaptic targets. We defined Aorta downstream interneurons as left/right homologous pairs in which at least one neuron received ≥3 synapses from Aorta sensory neurons, and mapped their output to downstream target groups (**Figure 5A**).

**Figure 5.**
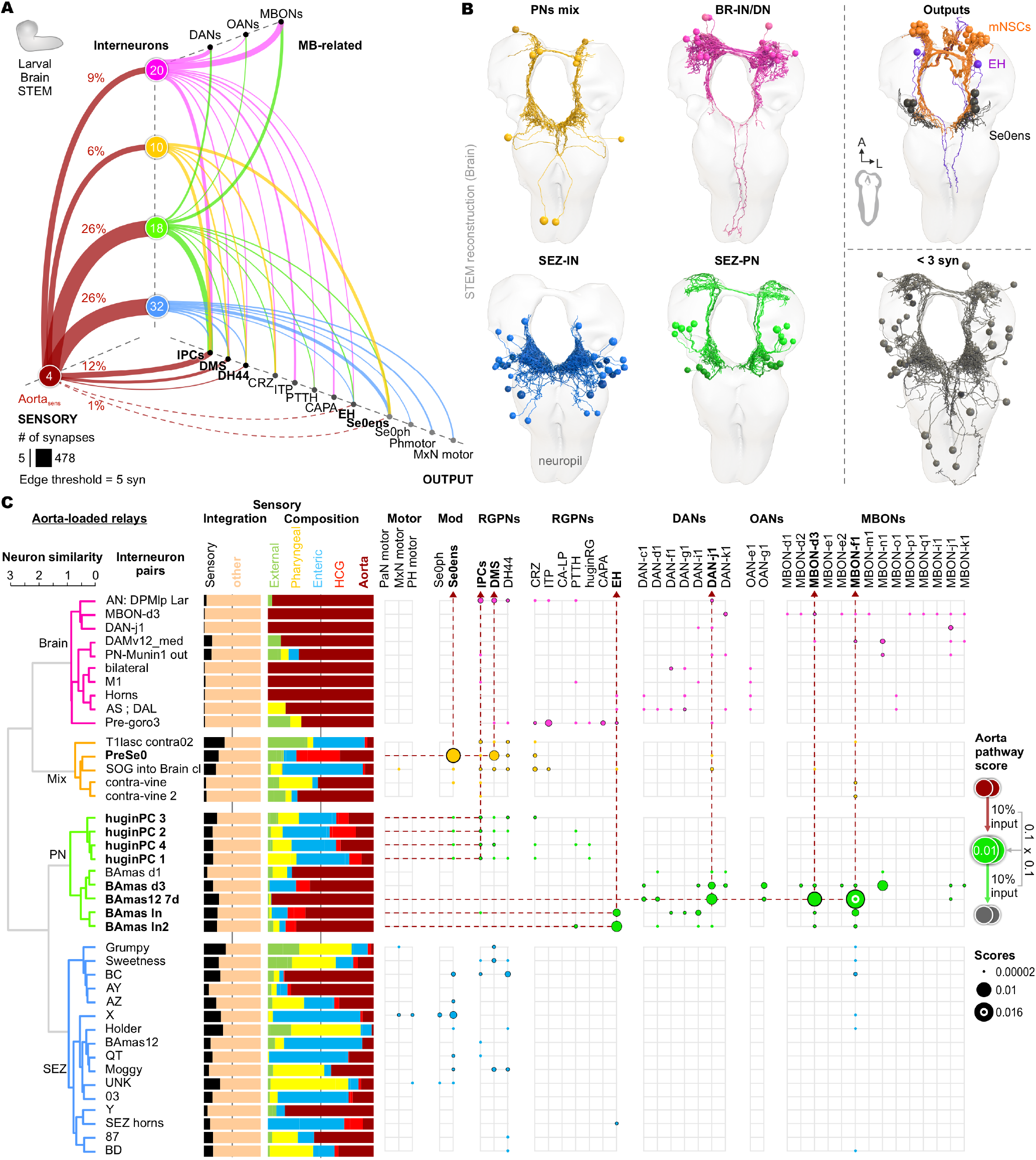
Aorta sensory neurons distribute hemolymph-associated input through morphologically distinct interneuron relays. **(A)** Beehive plot showing the complete direct connectivity of aorta sensory neurons to nervous-system output neurons and downstream interneuron pairs. Interneurons were defined as aorta downstream neurons when at least one neuron of a left/right homologous pair received ≥3 synapses from aorta sensory neurons. Connections from these interneuron pairs to mushroom body-related input and output neurons are also shown. Edges are displayed with a threshold of ≥5 synapses. Percentages indicate the fraction of the total outgoing synaptic budget of aorta sensory neurons. Interneurons are grouped by morphological similarity, as shown in C. **(B)** EM reconstructions of morphologically defined interneuron groups, direct output targets of aorta sensory neurons, and neurons below the aorta-downstream threshold. **(C)** Multilevel analysis of aorta downstream interneuron pairs. From left to right: morphology-based similarity calculated by NBLAST and shown as a dendrogram; corresponding interneuron-pair names; normalized total input of each pair, separating sensory input from other input; sensory composition of each pair, shown as stacked bar plots normalized to total sensory input and ordered according to the sensory-stage model from external to pharyngeal, enteric, HCG, and aorta input; and aorta-loaded relay analysis. In the relay matrix, dots indicate the presence of a relay path from Aorta sensory neurons to an interneuron pair to a downstream target group, and dot size represents the corresponding relay score. Relay-score calculation and dot-size scaling are shown on the right.

These interneuron pairs occupied distinct anatomical domains and clustered into four morphological classes: SEZ-local interneurons, SEZ-to-brain projection neurons, mixed ascending/descending neurons, and brain-local interneurons (**Figure 5B,C**). For each pair, we quantified the direct sensory input and normalized its composition by modality and origin, separating external, pharyngeal, enteric, HCG, and Aorta input (**Figure 5C**). This revealed that Aorta downstream interneurons are not a uniform relay population. Instead, the four classes differ in sensory load, sensory composition, and downstream target bias. SEZ-local interneurons received Aorta sensory input together with prominent gustatory and enteric sensory input. They formed multiple relay paths to PI neurosecretory cells and serotonergic output neurons, with more limited access to the feeding motor system. Mixed ascending/descending neurons showed the strongest relay to serotonergic output neurons, together with additional paths to PI and CRZ neurons. Brain-local interneurons received relatively little sensory input overall, but the sensory fraction they did receive was often dominated by Aorta input. This group formed many relay paths to DANs, OANs, and MBONs, as well as a smaller number of strongly combinatorial relays to PI and PL neurosecretory targets.

Within this overall architecture, several convergence motifs stood out. The SEZ-to-brain projection neuron class formed the most structured projection cluster and separated into two major groups: huginPC neurons and BAmas neurons. These groups define parallel channels from the SEZ toward higher-order output systems. HuginPC neurons link Aorta sensory input to endocrine targets, consistent with the previously described hugin circuit (Schlegel et al. 2016), whereas BAmas neurons provide a route from Aorta sensory input toward mushroom body-related circuitry (**Figure 5C**).

Additional motifs were organized around specific downstream targets. Interneurons relaying to Se0ens serotonergic output neurons included Pre-Se0 neurons, whose somata lie in the protocerebrum but whose output reaches Se0ens in the SEZ. The BC pair represented a particularly Aorta-loaded relay: its sensory input was dominated by Aorta sensory neurons, and its strongest relay path was directed to DH44 neurons, with additional relays to serotonergic output neurons and IPCs.

Together, this relay architecture shows that Aorta sensory neurons distribute hemolymph-associated information through multiple anatomically distinct interneuron classes. These relays connect internal state not only to neuroendocrine and serotonergic systems, but also to dopaminergic, octopaminergic, and mushroom body output pathways.

### Aorta sensory neurons connect neuroendocrine and mushroom body memory circuits

Among the major Aorta-associated projection neuron classes, Hugin PCs and BAmas define two parallel routes for Aorta-derived information (**Figure 6A**). Hugin PCs receive direct input from Aorta sensory neurons, but their strongest sensory input comes from enteric mechanosensory neurons. In contrast, BAmas neurons are dominated by Aorta sensory input, which accounts for most of their total sensory input (**Figure 6B**). Thus, Hugin PCs provide a route toward PI neuroendocrine targets, whereas BAmas neurons form a prominent projection-neuron relay from Aorta sensory neurons toward mushroom body-related circuitry (**Figure 6C**). At the group level, both projection-neuron classes are embedded in the broader Aorta downstream interneuron network, suggesting that Aorta-derived information may undergo network-level integration rather than being relayed through isolated feedforward paths.

**Figure 6.**
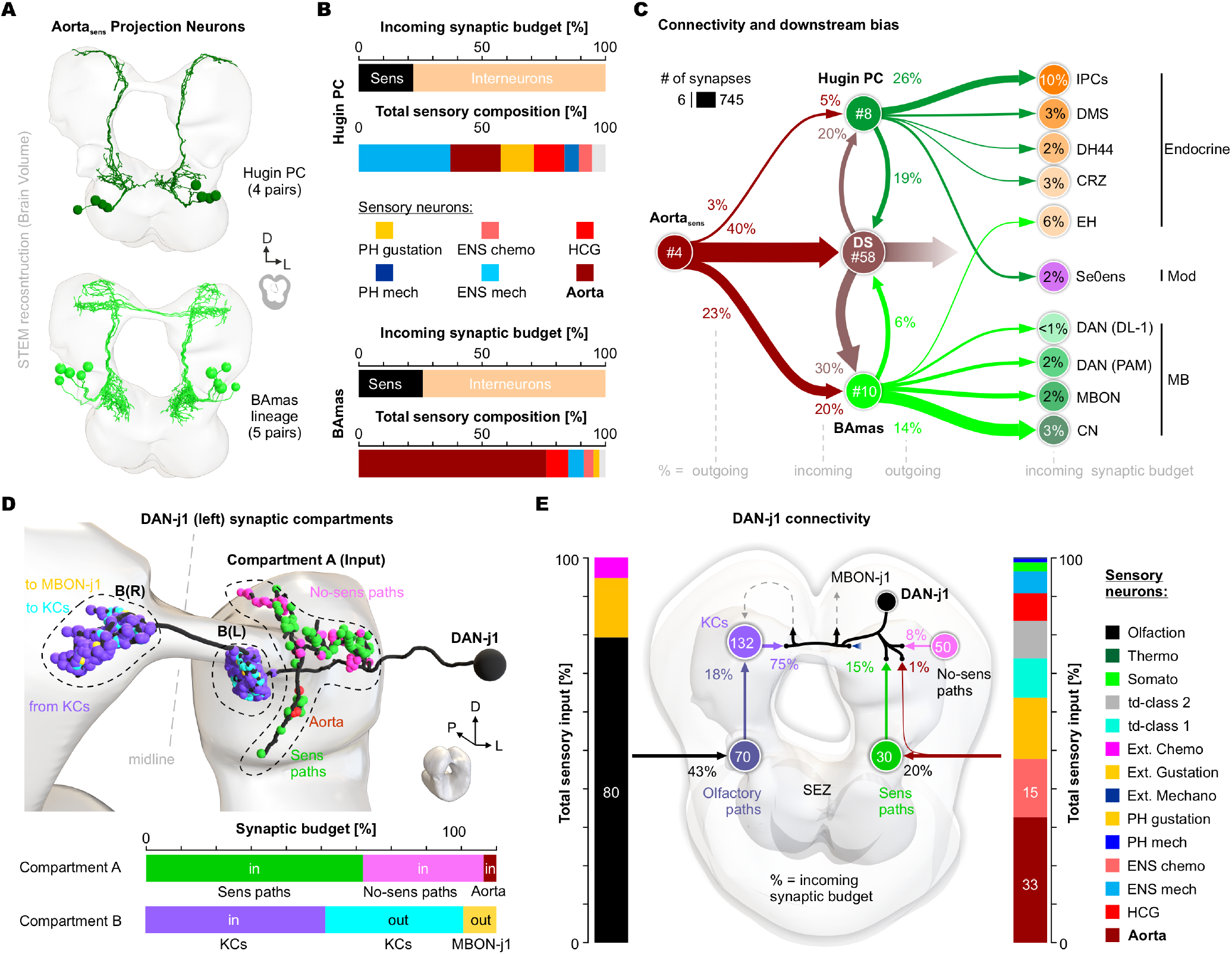
BAmas neurons are prominent aorta-associated projection neurons and provide relay pathways to mushroom body neurons. **(A)** STEM reconstructions of Hugin PCs and five BAmas lineage pairs belonging to morphological group 2 (SEZ projection neurons). This panel summarizes the major aorta-associated projection neurons. **(B)** Comparison of sensory input relative to total synaptic input in Hugin PCs and BAmas neurons. The sensory composition of each group is shown below. Notably, 75% of the total sensory input to BAmas neurons originates from aorta sensory neurons. **(C)** Connectivity of Hugin PCs and BAmas neurons within the aorta-associated relay network. BAmas neurons are the strongest postsynaptic target class of aorta sensory neurons among these projection-neuron pathways and can be assigned, at least in part, to mushroom body-related circuits. In contrast, Hugin PCs represent an endocrine/pars intercerebralis-associated route for aorta-derived information. **(D)** Compartmental organization of DAN-j1. DAN-j1 can be subdivided into distinct synaptic compartments: an input compartment (Compartment A), located directly distal to the primary neurite, which receives sensory-pathway input and input from interneurons lacking sensory input; and a mushroom body compartment B, B(R) and B(L), which is both input and output compartment for Kenyon cells and provides output to MBON-j1. **(E)** Complete connectivity of DAN-j1. On the left side of the brain, olfactory pathways to DAN-j1 are shown, including interneuron-mediated pathways and Kenyon cell input. Kenyon cell input accounts for 75% of all synaptic input to DAN-j1. On the right side, all interneuron pathways to DAN-j1 are shown and compared according to whether they carry sensory input or not. Aorta sensory neurons are the dominant sensory class within the sensory input paths. Percentages indicate the fraction of the total incoming synaptic budget contributed by the respective neurons.

We next examined how this internal sensory information reaches DAN-j1, a mushroom body dopaminergic neuron (Modi et al. 2020; Lin 2023; Truman et al. 2023). This analysis included the complete upstream of DAN-j1 and distinguished interneuron paths with sensory input from those without detected sensory input (**Figure 6D,E**). DAN-j1 showed two main synaptic compartments. A proximal compartment, located directly distal to the primary neurite, received broad interneuronal input. Within the sensory-bearing paths to this compartment, which include BAmas neurons, Aorta-derived input was the dominant sensory component. A second, more distal mushroom body compartment received Kenyon cell input and contained DAN-j1 output synapses onto MBON-j1 and Kenyon cells. Kenyon cell input accounted for 75% of all synaptic input to DAN-j1, indicating that the major drive remains mushroom body-derived, while internal sensory information reaches the same neuron through a distinct upstream compartment.

This organization places Aorta-derived input next to the much larger olfactory and mushroom body input paths, but at a different anatomical site. Olfactory information reaches mushroom body circuits through the canonical olfactory pathway and Kenyon cell input, whereas Aorta-derived internal-state information reaches DAN-j1 through sensory-driven interneuron relays. The convergence of these paths onto the same dopaminergic neuron suggests a circuit logic in which external food cues and internal metabolic state can be compared or combined during value updating (**Figure 6E**).

In this view, Aorta sensory neurons may report the internal metabolic consequences of food after ingestion, while olfactory and other external sensory pathways provide information about food before ingestion. This organization fits a five-stage feeding model in which external appraisal, oral sampling, swallowing, digestion, and hemolymph-based internal confirmation occur over different timescales. Aorta sensory input would therefore contribute to a late internal confirmation signal that updates future food valuation and action selection. More broadly, this links serotonin-associated motor control during swallowing with dopamine-associated updating of nutrient value, a relationship summarized in the model in (**Figure 7**).

**Figure 7.**
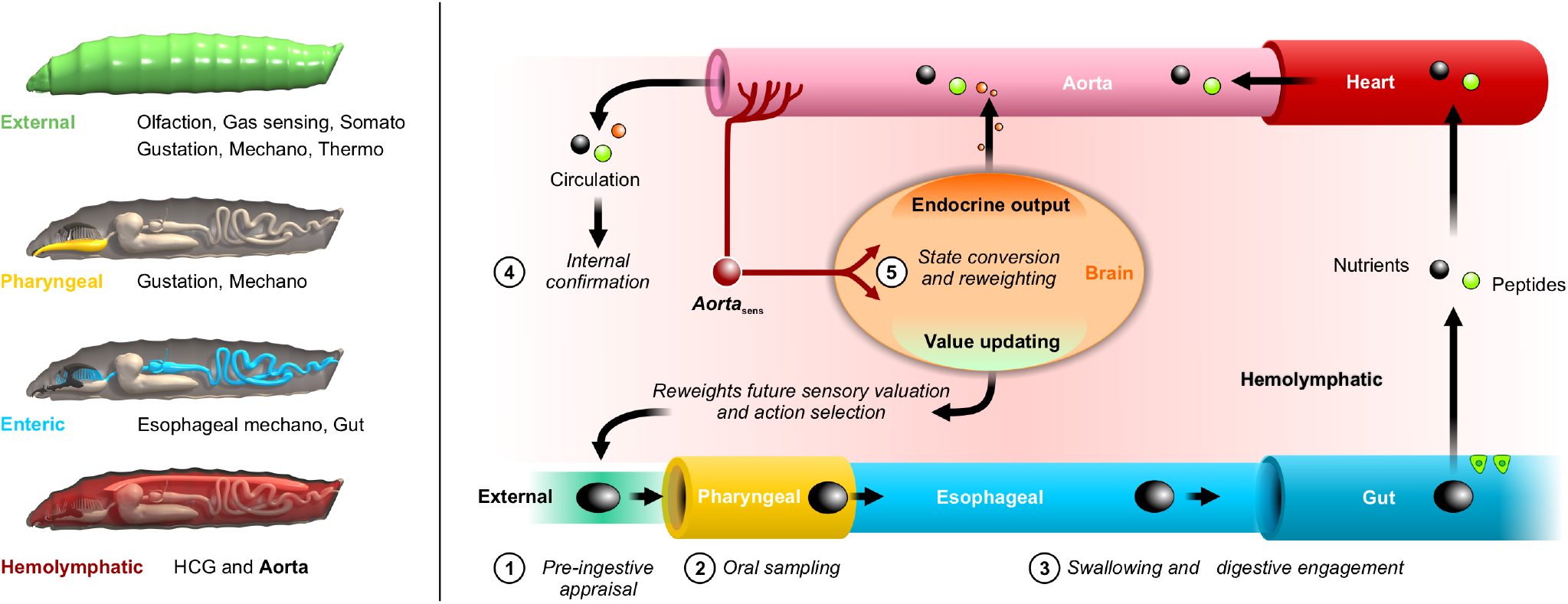
A five-stage multi-temporal feeding model links sensory input to metabolic state updating and future food valuation. **Left:** Overview of the major sensory regions of the *Drosophila* larva, organized along the feeding axis into external, pharyngeal, enteric, and hemolymphatic domains. The corresponding sensory modalities are indicated for each region, illustrating the anatomical distribution of sensory inputs across successive stages of food evaluation and ingestion. **Right:** Schematic of the five-stage feeding model. In **Stage 1**, food is evaluated by **pre-ingestive appraisal** through external sensory cues. In **Stage 2**, food is assessed by **oral sampling**. In **Stage 3, swallowing and digestive engagement** transfer ingested material into the gut. In **Stage 4**, food is broken down into nutrients that enter the hemolymph, while gut-derived peptides are additionally released into the circulation according to food content (these can also derive from other peripheral organs, such as the fat body). These signals are transported by the heart through the aorta to the aortic funnel, where the dendrites of aorta sensory neurons are located. Detection by aorta sensory neurons provides **internal confirmation** of nutrient availability. In **Stage 5**, this information is integrated in the brain to drive **state conversion, neuroendocrine output**, and **value updating**. These processes reweight future sensory valuation and action selection, thereby feeding back onto Stage 1 and closing the loop between feeding experience, internal metabolic state, and future food choice.

## DISCUSSION

We have identified the Aorta sensory neurons as a hemolymph-facing interoceptive component that links the peripheral circulation to central endocrine, feeding, and memory-related circuits. Their organisation differs fundamentally from brain-resident sensing mechanisms, where neurons or glia inside the CNS respond to peripheral signals that must first cross, or be carried across, the blood-brain barrier. Their dendrites are embedded in the wall of the aortic funnel and exposed to the hemolymph, while their axons enter the brain and synapse onto a defined set of central targets. They therefore provide a direct anatomical route through which circulating internal-state information can enter the nervous system with cellular and synaptic specificity.

### Peripheral interoceptors that synapse directly to the brain neurosecretory cells

A central feature of this organisation is the close coupling between neurosecretory release and sensory access at the aorta. Neurosecretory cells release into the aortic compartment, placing their peptides into the same circulating space sampled by Aorta sensory dendrites. This creates an anatomical loop: central neurosecretory cells release into the aorta, Aorta sensory neurons sample the aortic hemolymph compartment, and the same sensory pathway projects back to neurosecretory circuits in the brain. The loop is, for now, anatomical rather than chemically defined, because we do not yet know which receptors the Aorta sensory neurons use to detect any of the released peptides. It does, however, define a substrate for direct coupling between endocrine release and internal-state sensing. The receptor repertoire of Aorta sensory neurons suggests that this coupling is more than a direct nutrient readout. They express Gr43a and respond to fructose, but also serotonin receptors and the CCHa2R-RA isoform. CCHa2 is a nutrient-responsive peptide released from gut and fat body that rises after feeding and acts centrally on the IPCs to promote DILP release (Sano et al. 2015; Ren et al. 2015). Its receptor in Aorta sensory neurons raises the possibility that CCHa2 also acts on the sensory arm of the circuit, allowing the same peptide to be read twice along the insulin-control pathway: peripherally at the sensor and centrally at the effector. In this view, Aorta sensory neurons are not simple nutrient detectors, but peptide-gated interoceptors that combine metabolite sensing through Gr43a with humoral and modulatory information about feeding state.

The Aorta sensory organ is not the only internal interface in this system. The HCG represents a second hemolymph-associated site, but one with a different anatomical and functional bias. We previously placed the HCG within an esophageal sensorimotor module linked to mechanosensation, serotonergic modulation, and ring muscle control (Schoofs et al. 2024). Our current data add that HCG sensory somata and dendrites are exposed to the hemolymph, while their connectivity points to a target bias distinct from Aorta sensory neurons. Thus, Aorta sensory and HCG pathways appear to represent parallel but non-equivalent interoceptive routes. IPCs and DMS neurons receive strong direct input from Aorta sensory neurons, whereas DH44 neurons are more strongly associated with HCG input and broader multisensory pathways. DH44 neurons also respond to nutritive sugars and amino acids and drive post-ingestive food selection ((Dus et al. 2015; Yang et al. 2018; Oh et al. 2019), suggesting that they integrate a composite representation of the feeding event. IPCs, in contrast, receive a more privileged Aorta-associated signal, well positioned to couple circulating metabolic state to insulin-like peptide regulation.

DMS occupies an intermediate position, receiving strong Aorta input but also prominent HCG input, and may therefore sit at a convergence point between aortic and esophageal internal-state pathways. Myosuppressins are classic inhibitors of visceral muscle: Dromyosuppressin stops crop contractions and dampens gut motility and heart rate in a dose-dependent, tissue-specific way (Duttlinger et al. 2002; Richer et al. 2000; McCormick and Nichols 1993). In this view, IPCs would couple aortic metabolic state to systemic insulin-like peptide output, DH44 would integrate broader post-ingestive and HCG-associated signals, and DMS would link these internal-state pathways to visceral motor and endocrine stabilization. Both Aorta sensory and HCG routes are further distributed through interneuron pathways, arguing against a single internal-state channel from “the hemolymph” to “the brain” and instead for multiple peripheral interfaces feeding into overlapping downstream networks.

### Aortic interoception to post-ingestive metabolic reward

A striking aspect of the Aorta sensory pathway is its access to mushroom body-related circuitry (Eichler et al. 2017; Eschbach et al. 2020; Saumweber et al. 2018; Eschbach and Zlatic 2020; Weber et al. 2025; Schleyer et al. 2020). The direct contacts onto DAN-j1 and MBON-d3 are sparse in absolute terms, but they provide a minimal anatomical interface between hemolymph-associated sensory input and the reinforcement/value-updating system. Olfactory and Kenyon cell pathways remain quantitatively much larger, but the importance is anatomical specificity: Aorta-derived internal-state information reaches memory-related neurons at defined synaptic sites. The relay analysis expands this point. Hugin PCs and BAmas define two major SEZ-to-brain projection routes. Hugin PCs receive Aorta input, but their sensory drive is mixed and dominated by enteric mechanosensory input, in keeping with their established role in feeding suppression, bitter avoidance, and neuroendocrine access (Schlegel et al. 2016; Hückesfeld et al. 2016). BAmas form the complementary route. Their sensory input is dominated by Aorta sensory neurons, and their downstream reach is biased toward DANs, MBONs, and convergence neurons of the mushroom body output layer. This makes BAmas the principal projection-neuron route through which the hemolymph-associated state can be converted into memory-related value-updating signals.

DAN-j1 also illustrates how such integration could be organized inside a single neuron. Its total input is dominated by Kenyon cells, consistent with its role in mushroom body circuitry. However, DAN-j1 also contains a proximal input compartment, directly distal to the primary neurite, that receives broad interneuronal input. Within the sensory-bearing paths to this compartment, including BAmas-containing paths, Aorta-derived input is the dominant sensory component. The Kenyon cell-associated compartment lies further downstream and also contains DAN-j1 output sites to MBON-j1 and Kenyon cells. This topographic organisation suggests that internal hemolymph-associated input and canonical mushroom body input converge onto the same dopaminergic neuron, but at different anatomical sites along its arbor. Aorta-derived signals may therefore modulate the context in which Kenyon cell activity is evaluated, rather than simply adding another sensory input to the Kenyon cell layer.

This provides a plausible circuit solution to a delayed credit-assignment problem in feeding (Richards and Lillicrap 2019). External sensory systems, especially olfaction and taste, evaluate food before and during ingestion, while pharyngeal and enteric systems monitor oral sampling, swallowing, and digestive engagement. Sweet taste is enough to build short-term appetitive memory, but stable long-term memory requires the actual nutrient value of the sugar, and that value reaches the dopaminergic system only after ingestion and metabolism (Burke and Waddell 2011; Fujita and Tanimura 2011; Dus et al. 2011; Huetteroth et al. 2015; Yamagata et al. 2015; Musso et al. 2015). The afferent that carries this delayed post-ingestive signal to the dopaminergic neurons has not been identified. The Aorta sensory neurons are a strong candidate. They are positioned after food has been processed into circulating nutrients and peripheral signals in the hemolymph, and through direct and relay access to dopaminergic and mushroom body output circuits they could update the value assigned to the cues that preceded the meal. In that sense, the Aorta sensory pathway is a candidate substrate for metabolic memory.

This interpretation fits the five-stage feeding model proposed here (**Figure 7**). Food is first evaluated by external sensory cues, then by oral sampling, swallowing, and digestive engagement. Only later do nutrients and peripheral peptide signals enter the hemolymph and reach the aortic funnel. Aorta sensory neurons are positioned at this late internal stage, where they can report the metabolic consequence of feeding back to the brain, with the HCG providing a partly parallel internal route at the esophageal interface. The model therefore separates immediate feeding control from delayed value updating. Serotonin-associated circuits participate in swallowing, motor state, and feeding-state transitions, whereas dopamine-associated mushroom body circuits update the learned value of food according to its internal consequences. Together, Aorta sensory and HCG pathways suggest that the larval brain receives internal-state information through multiple peripheral checkpoints arranged along the feeding axis.

A broader implication is that gut-to-brain and circulation-to-brain pathways in invertebrates may carry teaching information, not only homeostatic information. In mammals, vagal and circumventricular pathways link visceral and circulating signals to hypothalamic, brainstem, and dopaminergic circuits, and recent work has emphasized that gut-derived signals can support reinforcement as well as satiation (Han et al. 2018). The Aorta sensory pathway is not anatomically homologous to these vertebrate systems. Yet they solve a comparable problem in a small brain: they convert delayed internal information into circuit-specific input to endocrine, motor-state, and memory networks. This makes Drosophila a tractable place to work out how an interoceptive signal becomes linked to reinforcement, endocrine control, and future choice.

Several limitations remain. The molecular ligands detected by Aorta sensory neurons and HCG sensory neurons are still largely unknown. Fructose responsiveness and expression of candidate receptors, including Gr43a, Gr28a, serotonin receptors, and CCHa2R-RA, provide entry points, but do not define the full sensory repertoire. In particular, whether CCHa2 activates, inhibits, or modulates the gain of Aorta sensory neurons remains to be tested. The proposed role in metabolic memory is based on anatomy, connectivity, calcium responses, and peptide-state phenotypes. Direct behavioral experiments will be required to test whether Aorta sensory neuron activity changes food learning, future food choice, and state-dependent action selection, for example in paradigms that dissociate post-ingestive value from taste. This will allow moving from a synapse-resolved map of hemolymph-facing sensory interfaces to a causal account of how internal metabolic state updates behavior.

## ACKNOWLEDGEMENTS

We thank Hamza Chihab, Chiyoung Choi, Chris Göschel, Estella Krämer, Hope Nitsche for their contributions with the calcium imaging and EM neuroanatomy, Jan Ache, Carlos Ribeiro and Meet Zandawala for exchange of information and discussions, Ingo Zinke for help with genetics, Alex Bates for help with BANC volume, and Tom Kazimiers for help with Igor dataset.

## AUTHOR CONTRIBUTIONS

Investigation: A.M.;

EM-dataset: A.C.;

EM-reconstruction: A.M., A.S., D.D.;

Computation/Coding: P.S., M.F., C.P.;

Analysis: A.M., A.S., P.S.

Visualization: A.M.

Supervision: M.J.P.

Writing – original draft: A.M., M.J.P.

## FUNDING

German Research Foundation PA 787/7-3 and 787/9-3 (MJP)

German Excellence Strategy EXC2151-390873048 (MJP)

## DATA AND MATERIAL AVAILABILITY

All data generated or analyzed during this study are included in the manuscript and supporting files. We used the whole animal STEM volume reported in Peale et al., (2024). Contact for dataset accession: A. Cardona.

## SUPPLEMENTARY MATERIALS

Figure S1

## MATERIAL AND METHODS

### Experimental model and subject details

#### Fly work

All larvae were maintained at 25°C under a 12 h light/dark cycle unless stated otherwise. For behavioral experiments, 4 h egg collections were performed on apple juice agar plates containing a load of yeast/water paste. After 48 h, larvae were transferred into vials containing standard cornmeal medium at a density of 60 larvae per vial. For other experiments, including functional imaging, electrophysiological recordings, and antibody stainings, 4 h egg collections were performed in vials containing standard cornmeal medium with a spot of yeast/water paste, and animals were subsequently kept for 4 days at 25°C. Larvae used for optogenetic stimulation were raised on fly food containing 150 μM all-trans retinal (Sigma-Aldrich, R2500) and kept under dark conditions.

#### Fly lines and genotypes

All larvae used for experiments were 96 ± 2 h old. The following *Drosophila melanogaster* lines were used.

**Driver lines:** *peb*-Gal4 (BDSC #80570), CCHa2R-RA-Gal4 (BDSC #84604).

**Effector/reporter lines:** UAS-Cam2.1 (BDSC #6901), UAS-Chrimson (BDSC #55135), UAS-GCaMP6f (BDSC #42747), UAS-GFP (BDSC #32184), UAS-GtACR1 (BDSC #92983), UAS-myrGFP (BDSC #32197).

### Method details

#### Dissection of semi-intact larvae

Feeding third-instar larvae were dissected in Petri dishes coated with a two-component silicone elastomer (Wacker Chemical Corporation, Elastosil RT 601). Larvae were pinned dorsal side up at the posterior and anterior ends using sharp-etched tungsten needles with a diameter of 40–60 μm. The larva was opened longitudinally along the dorsal midline, and the cuticle was subsequently cut transversely below the cephalopharyngeal skeleton using micro scissors (Fine Science Tools, 15000-08). Internal organs, including fat body, trachea, and salivary glands, were removed, whereas the CNS, cephalopharyngeal skeleton, associated pharyngeal nerves, and digestive tract up to the anterior midgut were retained. This standard preparation of the *Drosophila* larva was used in all experiments involving larval dissection and is referred to here as the semi-intact preparation.

#### Immunohistochemistry

Dissected larval or adult brains were fixed for 1 h in 4% paraformaldehyde in 1× phosphate-buffered saline (PBS), rinsed three times for 20 min in 1% PBS-T (1% Triton X-100 in 1× PBS), and blocked for 2 h in 1% PBS-T containing 5% normal goat serum (ThermoFisher). Primary antibodies were added to the blocking solution at the concentrations listed below, and samples were rotated for two nights at 4°C. On the third day, primary antibodies were removed, and larval brains were washed three times for 20 min in 1% PBS-T and additionally blocked for 30 min in 1% PBS-T containing 5% normal goat serum. Secondary antibodies were then applied, and samples were rotated for two nights at 4°C. After three washes for 20 min in 1% PBS-T, brains were dehydrated and cleared through an ethanol-xylene series and mounted in DPX Mountant (Sigma-Aldrich).

Imaging was performed on a Zeiss LSM 780 confocal microscope using either an LCI Plan-Neofluar 25×/0.8 Imm Korr DIC M27 objective or a Plan-Apochromat 63×/1.4 Imm DIC oil objective.

For antibody staining of driver > GFP or driver > myrGFP samples, chicken anti-GFP was used as primary antibody (1:500; Abcam, ab13970), and goat anti-chicken Alexa Fluor 488 was used as secondary antibody (1:500; Invitrogen, A-11039). For pericardin/EC11 staining, mouse anti-EC11 was used as primary antibody (1:1000; DSHB, AB 528431), and goat anti-mouse Alexa Fluor 633 was used as secondary antibody (1:500; Invitrogen, A-21052).

For neuropeptide stainings, guinea pig anti-Dilp2 was used as primary antibody (1:500; Coring; antigen sequence DMKALREYCSVVRN), and goat anti-guinea pig Alexa Fluor 633 was used as secondary antibody (1:500; Invitrogen, A-21105). Rabbit anti-DH44 was used as primary antibody (1:500; Jan Veenstra), and goat anti-rabbit Alexa Fluor 633 was used as secondary antibody (1:500; Invitrogen, A-21071). Guinea pig anti-DMS was used as primary antibody (1:1000; Jan Veenstra), and secondary antibodies were goat anti-rabbit Alexa Fluor 633 (1:500; Invitrogen, A-21071) and goat anti-guinea pig Alexa Fluor 633 (1:500; Invitrogen, A-21105).

For background staining, mouse anti-Brp was used as primary antibody (1:500; DSHB, AB2314866), and goat anti-mouse Alexa Fluor 633 was used as secondary antibody (1:500; Invitrogen, A-21052). For occasional F-actin staining, conjugated fluorescent Phalloidin-TRITC was used (1:1000; Sigma-Aldrich, P1951).

#### Live calcium imaging of aorta sensory neurons

For functional imaging of aorta sensory neurons, GCaMP6f was expressed in sensory neurons using the pan-sensory driver *pebbled*-Gal4. Aorta sensory neurons were volumetrically imaged live in dissected larvae in Ringer’s solution. Glucose, trehalose, and fructose were tested as stimuli, and Ringer’s solution without sugar served as control. For each animal, solutions were applied sequentially in the following order: 0 mM sugar, 10 mM sugar, 0 mM sugar, and 50 mM sugar. Fluorescence changes were quantified from regions of interest corresponding to individual aorta sensory neuron somata and plotted as calcium-response traces. Imaging was performed at a sampling rate of 0.2 Hz. Statistical comparisons between control and stimulus conditions were performed using the Mann–Whitney U test.

#### Optogenetic manipulation and quantitative peptide immunostaining

For bidirectional optogenetic manipulation, CCHa2R-RA-Gal4 was used to drive either UAS-CsChrimson or UAS-GtACR1 in aorta sensory neurons. Larvae were raised on instant fly food with or without 150 μM all-trans retinal and used at 96 ± 2 h after egg laying. For long-term manipulation, larvae were exposed for 12 h to cyclic light stimulation with 15-min light-on/light-off intervals, using 590 nm light for CsChrimson-mediated activation and 520 nm light for GtACR1-mediated inhibition. Larvae were then dissected and immunostained for Dilp2. Dilp2 immunofluorescence was quantified and compared between control and experimental conditions.

For time-resolved peptide measurements after activation, CCHa2R-RA-Gal4 > UAS-CsChrimson larvae were transferred at 96 ± 2 h after egg laying to Petri dishes containing filter paper moistened with 1× PBS and kept without food. Animals were exposed either to darkness or continuous 590 nm light and collected after 0, 1, 2, and 4 h. Larvae were dissected immediately after collection and immunostained for Dilp2, DH44, or DMS. Matched control and experimental samples were processed in parallel and imaged on a Zeiss LSM 780 confocal microscope using identical acquisition settings within each experiment.

Mean fluorescence intensity was measured in the corresponding neurosecretory cell bodies. The primary readout was the difference in peptide immunofluorescence between non-activated controls and animals with optogenetically activated aorta sensory neurons. Relative fluorescence intensity was calculated by dividing individual fluorescence values from light-stimulated animals by the mean fluorescence intensity of the corresponding dark-control group at the same time point. Fold changes were log-transformed for plotting. Peptide immunofluorescence was used as an indirect proxy for intracellular peptide content, peptide depletion, and putative release, but not as a direct measurement of secretion. Statistical comparisons were performed using the Mann–Whitney U test.

#### Quantification and statistical analysis

All ImageJ and Spike2 scripts used to analyze physiological datasets are available at https://github.com/Pankratz-Lab.

All statistical analyses were carried out in SigmaPlot 12.0 or GraphPad Prism 9.0. All statistical tests, replicate numbers, and statistical significance values for represented data are reported in the corresponding figures, figure legends, and relevant Method details sections. In all boxplots shown in figures and supplementary figures, the solid line depicts the median, and the upper and lower boundaries of the box depict the first and third quartiles of the dataset, respectively. Whiskers indicate the 5% and 95% confidence levels. Individual data points are shown as circles. All measurements were taken from distinct samples.

### EM reconstruction

#### Whole-animal volume

Neuron reconstruction was performed on a scanning transmission electron microscopy volume of a whole first-instar larva. Technical details of volume generation are described separately in Peale et al. 2024. Reconstructions were made in a modified version of CATMAID (http://www.catmaid.org). To reconstruct a neuron, a specific neurite was identified in a section of the STEM dataset, and a three-dimensional neuronal skeleton, including synaptic active zones and synaptic partners, was manually generated.

All enteric neurons, Se0 neurons, and specific pharyngeal neurons were identified by reconstructing all axons passing through the frontal nerve junction and originating either in the CNS or the enteric nervous system. All peripheral projections of the neurosecretory cells of the pars intercerebralis and pars lateralis were reconstructed. Neurons were reconstructed to completion, with 100% tracing and at least 95% review. All tissues innervated by aorta sensory neurons, including the aorta, heart, ring gland, and aorto-esophageal muscles, were reconstructed.

#### Brain volume

Neuron reconstruction was performed on an ssTEM volume of a 6-h-old first-instar larva (Ohyama et al. 2015). For the sensory neurons included here, previously published data were used (Berck et al. 2016; Miroschnikow et al. 2018; Ohyama et al. 2015; Schlegel et al. 2016; Hückesfeld et al. 2021). For interneurons and output neurons included here, previously published data were used (Berck et al. 2016; Miroschnikow et al. 2018; Schlegel et al. 2016; Hückesfeld et al. 2021; Schoofs et al. 2024; Eichler et al. 2017; Winding et al. 2023; Eschbach et al. 2020; Ohyama et al. 2015). All synaptic upstream and downstream partners of aorta sensory neurons were reconstructed to completion using a synaptic threshold of three synapses.

#### Relay-score calculation

To quantify indirect sensory relay strength from defined sensory neuron groups to downstream target groups, we calculated a relay score for each interneuron pair. Sensory neurons were treated as groups, and interneurons were analyzed as left/right homologous pairs. For each interneuron pair, we first determined the fraction of its total synaptic input originating from a given sensory group. This sensory input fraction was then combined with the fraction of total target-group input provided by the same interneuron pair.

Targets were analyzed exclusively as defined target groups. For each target group, synaptic output from an interneuron pair and total target input were summed across all neurons belonging to that group. For aorta sensory neurons, the sensory group corresponds to the complete aorta sensory neuron population. The resulting relay score estimates how strongly an interneuron pair transmits input from a defined sensory group to a downstream target group, while accounting for both the sensory input received by the interneuron pair and the relative contribution of that pair to the target group’s total synaptic input.

#### Sensory-stage classification

For sensory-stage analysis, 15 sensory classes were grouped according to a feeding-related temporal model. External sensory inputs were assigned to Stage 1 and included olfaction, somatosensation, thermosensation, td-class2, td-class1, external chemosensation, external gustation, and external mechanosensation. Pharyngeal sensory inputs were assigned to Stage 2 and included pharyngeal gustation and pharyngeal mechanosensation. Enteric sensory inputs were assigned to Stage 3 and included ENS mechanosensation and ENS chemosensation. Hemolymphatic/internal sensory inputs were assigned to Stage 4 and included HCG1+2, HCG7+8, and aorta sensory neurons.

For each interneuron pair or target group, synaptic input from each sensory class was summed within the corresponding sensory stage. Sensory-stage fractions were then normalized to the total sensory input received by that interneuron pair or target group and displayed as stacked bar plots.

## Supplemental figures

**Figure S1.**
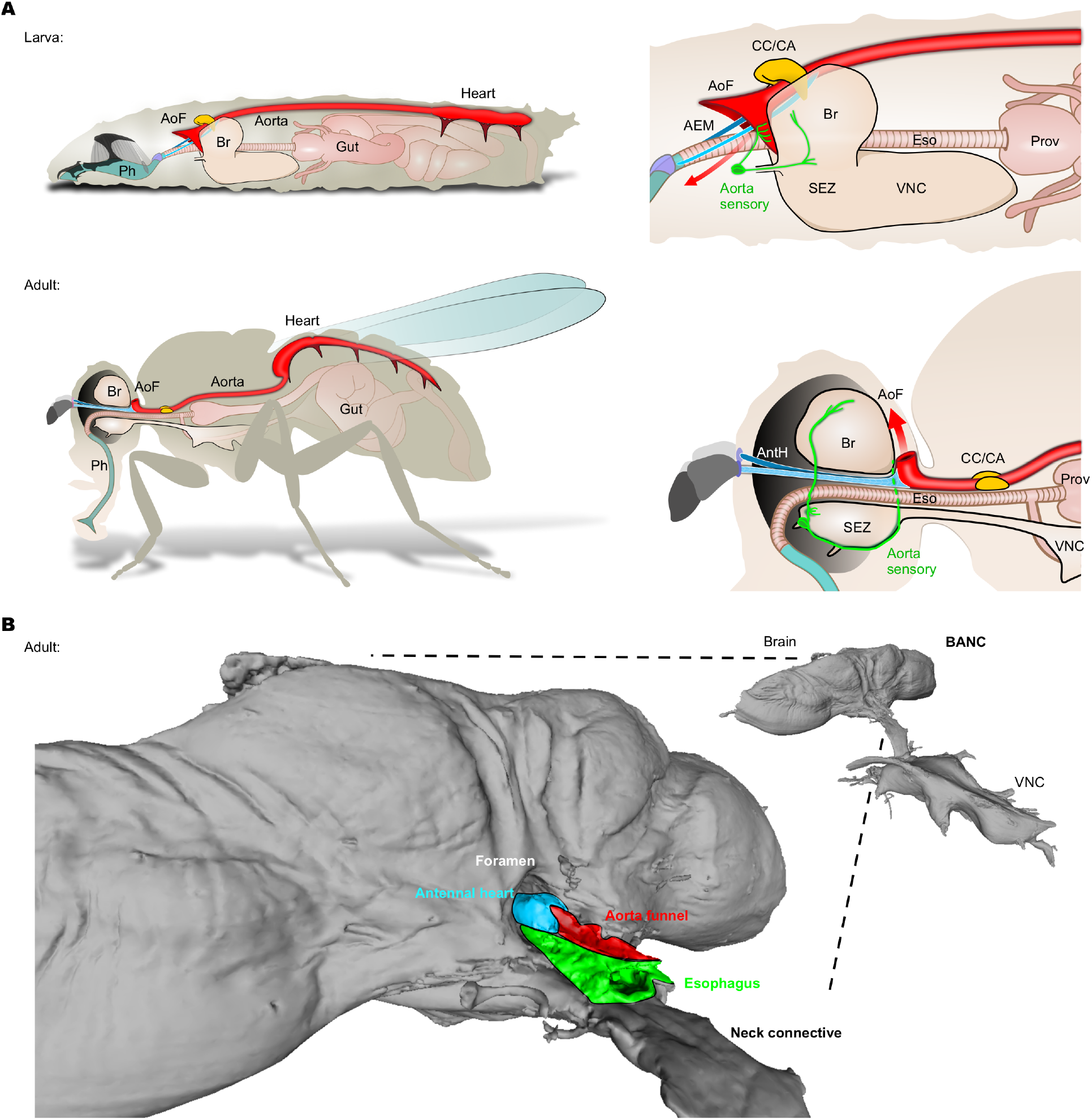
Reconstructed CNS-ENS connectome of *Drosophila* larva. (**A**) A schematic comparison of the circulatory system in larva and adult, based on current study and those of Demerc. **(B)** The BANC volume (Bates et al, 2026) contains a portion of the tissues that go through the foramen. Note that in most preparations for imaging, the aorta and the associated organs are removed or ripped off.

